# Susceptibility of bovine respiratory and mammary epithelial cells to avian and mammalian derived clade 2.3.4.4b H5N1 highly pathogenic avian influenza viruses

**DOI:** 10.1101/2025.01.09.632235

**Authors:** Victoria Meliopoulos, Sean Cherry, Maria Smith, Bridgett Sharp, Pamela H. Brigleb, Ericka Roubidoux, Brandi Livingston, Dorothea R. Morris, Tyler Ripperger, Pari Baker, Sara Gragg, Kiril Dimitrov, Stephanie Langel, Stacey Schultz-Cherry

## Abstract

Zoonotic transmission of avian influenza viruses into mammals is relatively rare due to anatomical differences in the respiratory tract between species. Recently, clade 2.3.4.4b highly pathogenic H5N1 avian influenza viruses were detected circulating in North American cattle. Sporadic transmission between cattle, humans, and other animals proximal to cattle or after consuming products from infected cattle has occurred, but thus far there is no evidence of human-to-human transmission. However, the virus has the potential to adapt to the mammalian respiratory tract with every transmission event that occurs, making it crucial to understand cellular and species tropism of the H5N1 2.3.4.4b viruses. We compared viral kinetics of clade 2.3.4.4b viruses isolated from birds and mammals in respiratory epithelial cells derived from cattle, human, swine, and ferret. We found that avian derived viruses could replicate in swine cells only, yet mammalian derived strains could replicate efficiently in all tracheal and nasal epithelial cells tested. Interestingly, only bovine mammary epithelial cells (MEC) and swine respiratory epithelial cells were permissive to both avian and mammalian derived strains, possibly due to increased sialic acid expression on bovine MEC compared to bovine tracheal epithelial cells (TEC). However, sialic acid expression differed between dairy and beef cows: TEC derived from a dairy cow had increased expression of α2,3 sialic acid receptors compared to TEC from a beef-dairy cow cross. This study highlights the ability of clade 2.3.4.4b H5N1 viruses derived from mammals but not wild birds to infect the respiratory epithelium of other mammalian hosts.

## Introduction

Influenza viruses cause acute respiratory infection in humans and other mammals. Influenza primarily targets epithelial cells of the respiratory tract; however certain influenza strains may display broader cellular tropism and can cause systemic infection sometimes resulting in a fatal outcome. The current outbreak highly pathogenic avian H5N1 clade 2.3.4.4b influenza viruses is one such example. Studies using animal models have shown these viruses have an extensive cellular tropism not limited to the respiratory tract (1, 2). In spring 2024, a clade 2.3.4.4b H5N1 influenza strain was detected circulating among dairy cattle (3). Surprisingly, the virus causes minimal respiratory disease (4, 5) but was replicating in an unexpected site: the mammary epithelium (6). However, how respiratory and mammary epithelial cells shape H5N1 influenza virus replication kinetics are unknown.

Primary respiratory epithelial cells can be cultured at the air-liquid interface and mimic the respiratory tract, developing tight junctions, mucins, and differentiation into a heterogenous population including goblet cells, basal cells, and ciliated cells. Cells isolated from different areas of the respiratory tract retain features of those areas, modeling the lung, the trachea, and even the nasal cavity (7). Importantly, these cultures can be generated from a variety of mammals allowing us to assess zoonotic potential of emerging influenza strains *ex vivo* (8, 9). These cells are a valuable tool considering the potential for emergence of an H5N1–seasonal influenza virus reassortant or viral adaptation to the mammalian respiratory tract.

Like the respiratory tract, the mammary gland has distinct populations of epithelial cells. Luminal and myoepithelial cells work in tandem to secrete milk and expel it from the duct (10). While it is surprising that high amounts of H5N1 are being detected in milk, viruses like human immunodeficiency virus (HIV), cytomegalovirus (CMV), and Zika virus can be transported to the mammary gland via the blood or lymphatic system resulting in viral shed into breast milk (11–13). A few studies have investigated influenza in the context of the mammary gland, showing intraductal inoculation of both ferrets and cattle resulted in shed of infectious virus in milk. Like respiratory epithelial cells, primary mammary epithelial cells can be differentiated in culture and can even be induced to secrete milk (14). In the context of virology, bovine mammary epithelial cells have been used to propagate viruses including influenza, and contain influenza-specific receptors (15, 16).

In these studies, we isolated and cultured primary differentiated respiratory and mammary epithelial cells from multiple mammalian species and assessed replication kinetics of clade 2.3.4.4b influenza viruses. We found that respiratory epithelial cells from all species tested could support replication of 2.3.4.4b viruses derived from mammals but none could support robust replication of wild bird isolates, except for swine cells. However, bovine mammary epithelial cells could support robust replication of most avian derived strains tested in addition to mammalian 2.3.4.4b strains, but not human seasonal H1 and H3 strains of influenza. This work gives insight to species tropism and possibly to our understanding of transmission dynamics of bovine H5N1 virus.

## Results

### Replication of mammalian derived 2.3.4.4b viruses in tracheal epithelial cells

Clade 2.3.4.4b H5N1 viruses can infect a wide range of hosts, and display tropism for cells beyond the respiratory tract (1, 2). We used primary respiratory epithelial cells from multiple mammalian species to explore the species tropism of clade 2.3.4.4b viruses. Previously we developed respiratory epithelial cells from swine (9) and ferrets (unpublished data), but since the current outbreak is occurring in cattle, we isolated tracheal epithelial cells (TEC) from lactating Holstein dairy cow (dairy) and a non-lactating, beef cow crossed with dairy (beef x dairy). Cells were cultured on transwell inserts at the air-liquid interface (ALI) until fully differentiated. Both bovine TEC cultures expressed high levels of the epithelial marker pan-keratin and exhibited hallmarks of differentiation such as mucus secretion and the presence of cilia (9, 17) (Figure 1).

**Figure 1.**
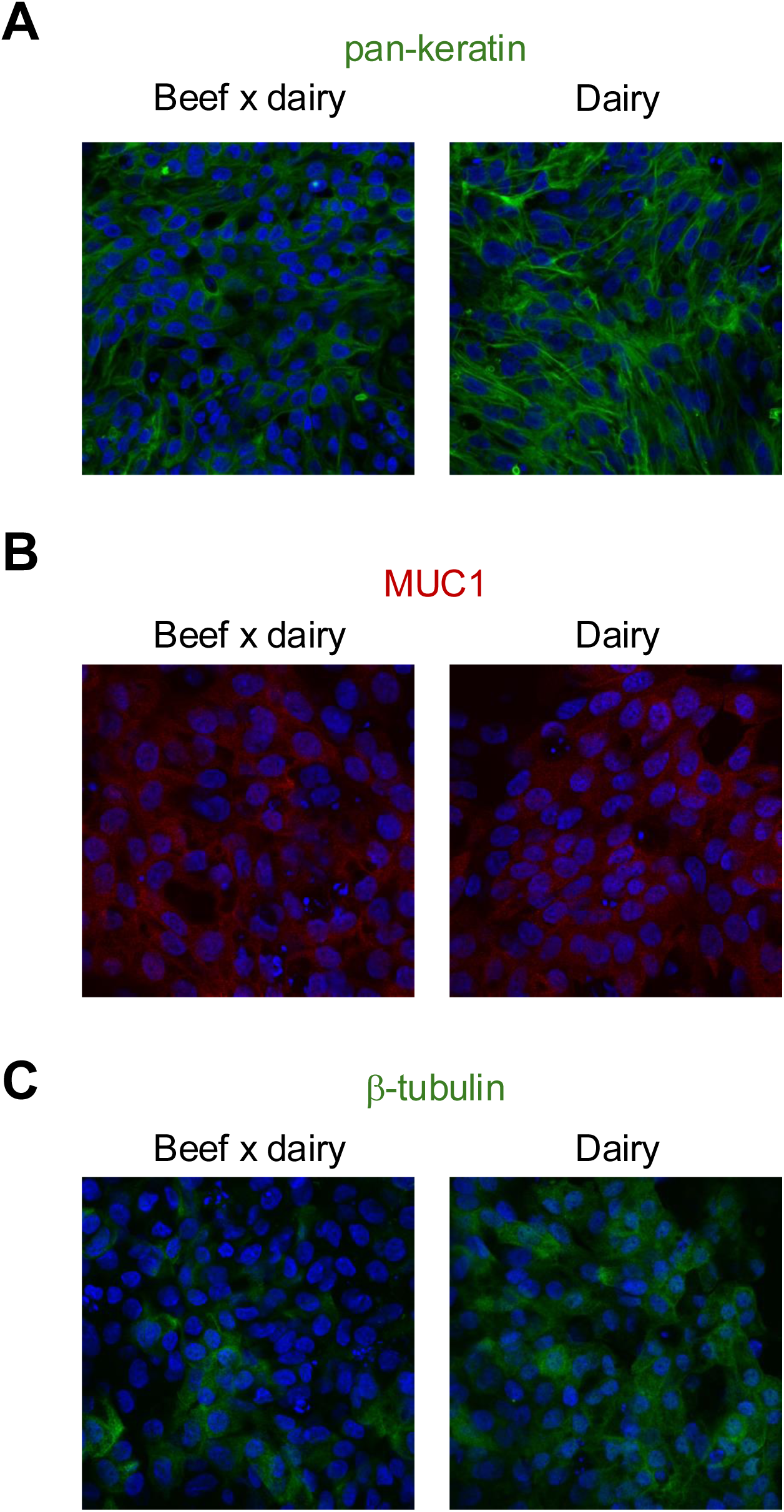
Bovine tracheal epithelial cells differentiate in culture. Tracheal epithelial cells (TEC) isolated from a beef x dairy and a dairy cow were cultured on transwell inserts until fully differentiated. Differentiated cells were fixed and immunostained for **(A)** pan-keratin (green), **(B)** MUC1 (red) and **(C)** β-tubulin (green). Cells were imaged by confocal microscopy. All samples were counterstained with DAPI (blue). Magnification is 40X. Data represents 2 independent experiments of n=2 per group.

Once we confirmed the viral isolates could replicate in MDCK cells (Figure S1), TEC were infected with mammalian derived clade 2.3.4.4b viruses from a cow [A/bovine/Ohio/B24OSU-439/2024; (bovine)] and a cat [A/feline/New Mexico/2024; (feline)] at a multiplicity of infection (MOI) of 0.5. Both strains replicated in tracheal epithelial cells, with the highest titers in swine TEC, followed by dairy, beef x dairy, and finally ferret TEC (Figure 2A-D). Interestingly, for the feline strain, the difference in viral titer was significantly different between swine TEC and both cow TEC (Figure 2D). Neither type of cow TEC could support replication of A/Switzerland/9715293/2013 (H3N2) or A/Michigan/45/2015 (H1N1), although the dairy TEC reached a peak titer of 10^5.6^ TCID_50_/mL at 24 hours post-infection (hpi) (Figure 2E). Overall, these data suggest that clade 2.3.4.4b viruses replicate efficiently in primary mammalian respiratory epithelial cells, particularly swine.

**Figure 2.**
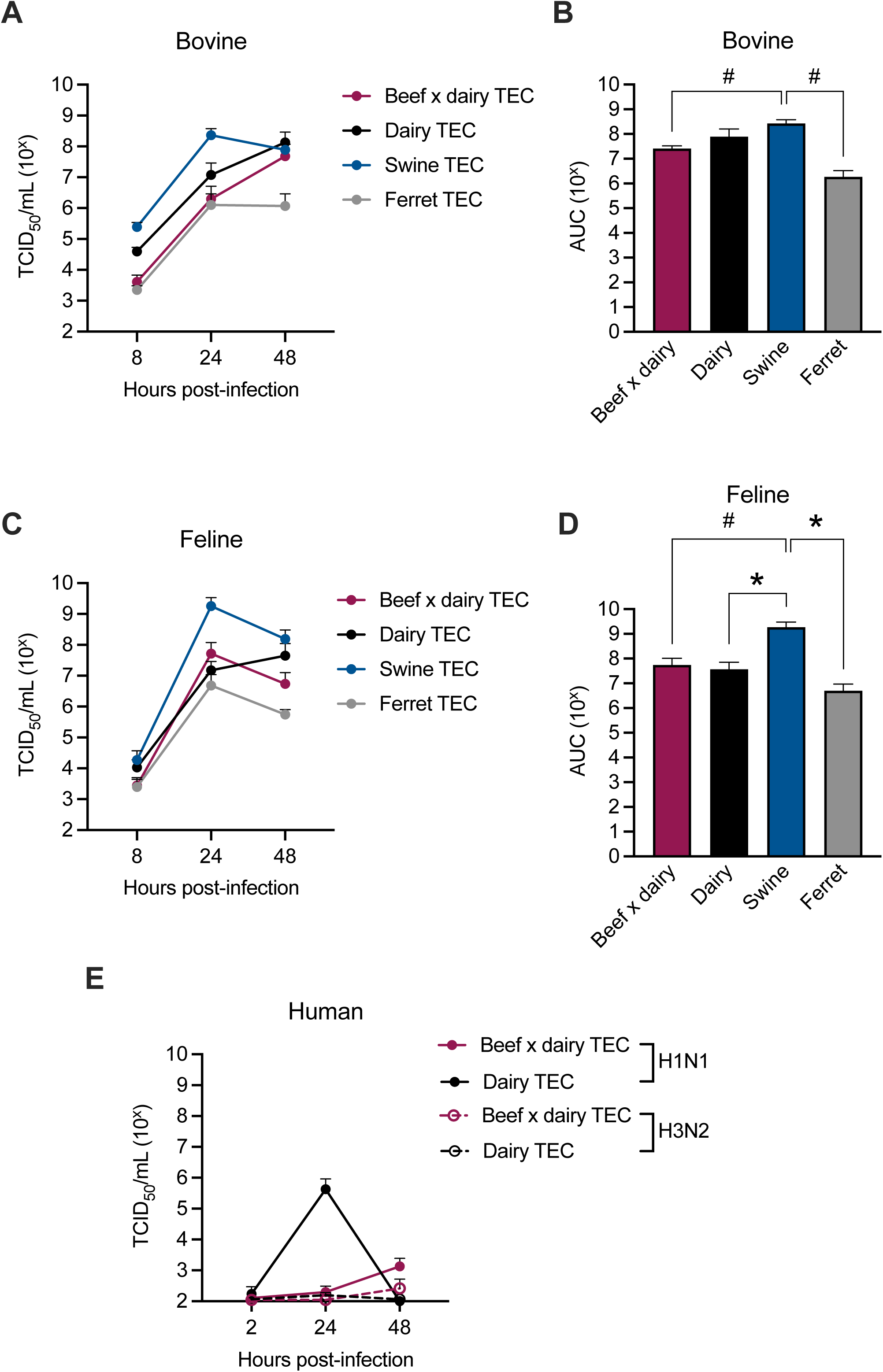
Mammalian derived clade 2.3.4.4b viruses replicate in tracheal epithelial cells isolated from multiple species. **(A)** Differentiated tracheal epithelial cells (TEC) from cows, swine, and ferret were infected with A/bovine/Ohio/B24OSU-439/2024 (H5N1) (bovine) at an MOI of 0.5. Apical washes were collected at the indicated time and presence of infectious virus was measured by TCID_50_ assay. **(B)** Area under the curve (AUC) of (A). Swine vs ferret *p*=0.0642; swine vs beef x dairy *p*=0.0992. **(C)** TEC were infected A/feline/New Mexico/2024 (H5N1) (feline) as in (A). **(D)** Area under the curve (AUC) of (C). Swine vs ferret *p*=0.0424; wine vs beef x dairy *p*=0.0504; swine vs dairy *p*=0.0475. **(E)** Cells were infected at an MOI of 0.5 with the seasonal human viruses A/Michigan/45/2015 (H1N1) or A/Switzerland/9715293/2013 (H3N2) as in (A). Significance of (B) and (D) was determined by one-way ANOVA with multiple comparisons, \**p*<0.05, ^#^p<0.10. Data represents 1 independent experiment of *n*=3-7/group. Error bars indicate standard error of the mean (SEM).

### Replication kinetics of mammalian derived 2.3.4.4b viruses in nasal epithelial cells

The upper respiratory tract, particularly the nasal cavity, is the primary site of influenza infection in mammals (18). To determine whether the mammalian derived 2.3.4.4b viruses could establish infection in nasal cells, we infected fully differentiated nasal epithelial cells (NEC) isolated from swine, ferret, and human. Unfortunately, we did not have access to bovine nasal cells. Both bovine and feline isolates replicated to high titers in all NECs tested. Like the TECs, swine NEC best supported replication of both strains, followed by human and ferret (Figure 3A-C). The bovine virus titers were significantly higher in the swine NEC compared to both ferret and human, however there was no significant difference in swine and human titers when infected with the feline virus (Figure 3C).

**Figure 3.**
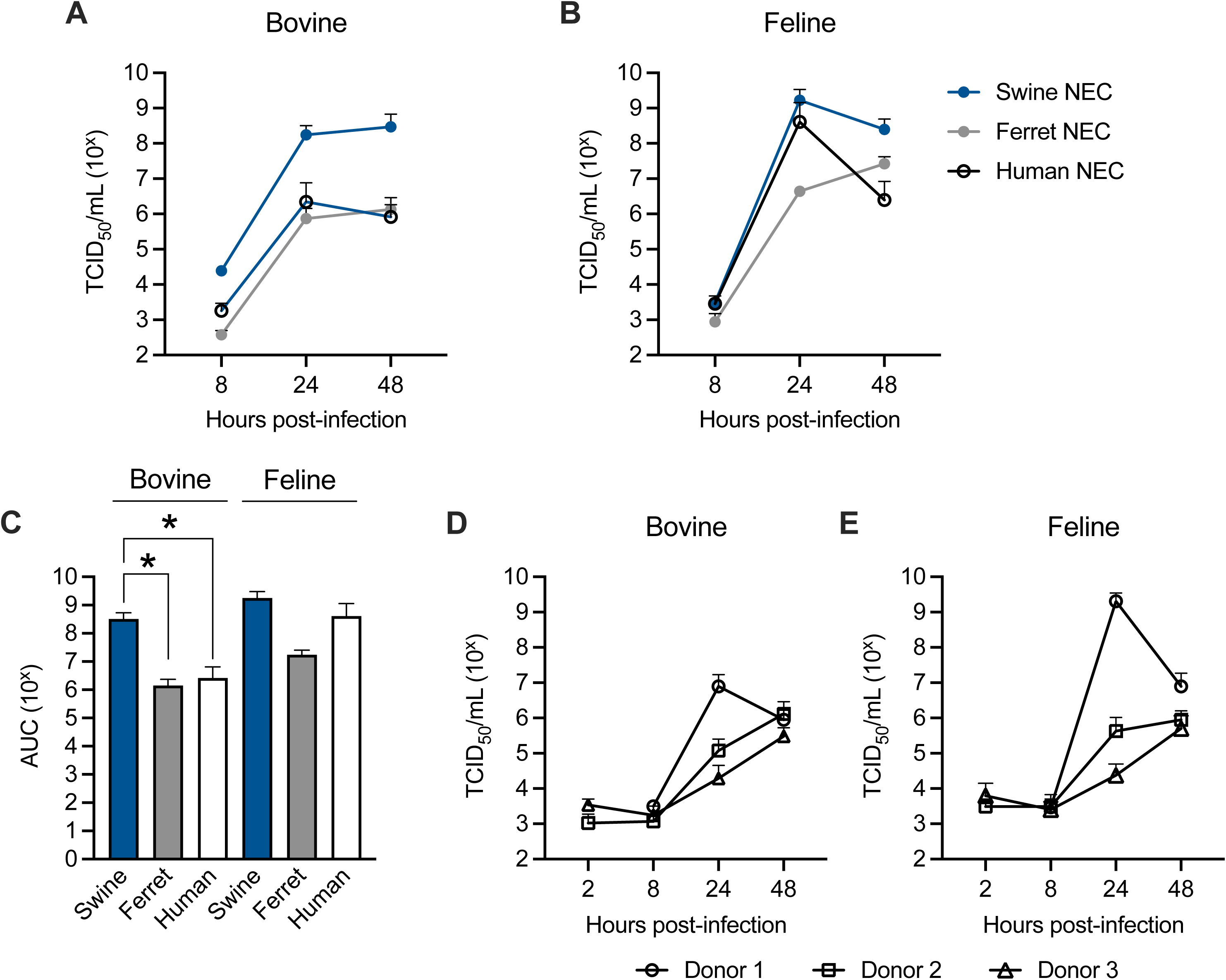
Mammalian derived clade 2.3.4.4b viruses replicate in nasal epithelial cells. Differentiated nasal epithelial cells isolated from swine, ferret, and human were infected with **(A)** A/bovine/Ohio/B24OSU-439/2024 (H5N1) (bovine) or **(B)** A/feline/New Mexico/2024 (H5N1) (feline) at an MOI 0.5. Apical supernatants were collected at the indicated timepoints and levels of infectious virus determined by TCID_50_ assay. Human NEC data represents 3 independent donors. **(C)** Area under the curve (AUC) of (A-B). Bovine: swine vs ferret *p*=0.0030; swine vs human *p*=0.0014. Significance was determined by one-way ANOVA with multiple comparisons; \**p*<0.05, ^#^*p*<0.10. **(D)** Viral kinetics in human nasal epithelial cells infected with A/bovine/Ohio/B24OSU-439/2024 (H5N1) (bovine) or **(E)** A/feline/New Mexico/2024 (H5N1) (feline) stratified by donor. Data represents 1 independent experiment of *n*=2-7/group (swine and ferret) or 2 independent experiments of *n*=3-7/group. Error bars indicate standard error of the mean (SEM).

To account for donor-to-donor variability, we used 3 independent human nasal cell donors. When the human data was stratified by donor, we saw the feline virus replicated to higher titers in one donor compared to the others, but both bovine and feline viruses were able to consistently replicate in human NEC (Figure 3D-E). Taken together, the bovine and feline isolates replicated in each respiratory cell type tested, especially swine. This may explain the recent identification of a clade 2.3.4.4b virus in swine in the U.S. (19) and highlights the need for increased surveillance.

### Avian derived clade 2.3.4.4b viruses do not replicate in primary mammalian respiratory cells

The infection of mammals with clade 2.3.4.4b viruses is likely due to spillover from wild birds. Thus, we asked if wild bird derived clade 2.3.4.4b viruses replicated in mammalian respiratory epithelial cells. Differentiated TEC and NEC cultures were infected with A/bald eagle/Florida/W22-134-OP/2022 (eagle/FL) at an MOI of 0.5. The avian virus replicated poorly in all TEC and NEC except those from swine, which had significantly higher viral burden than all other species tested (Figure 4C).

**Figure 4.**
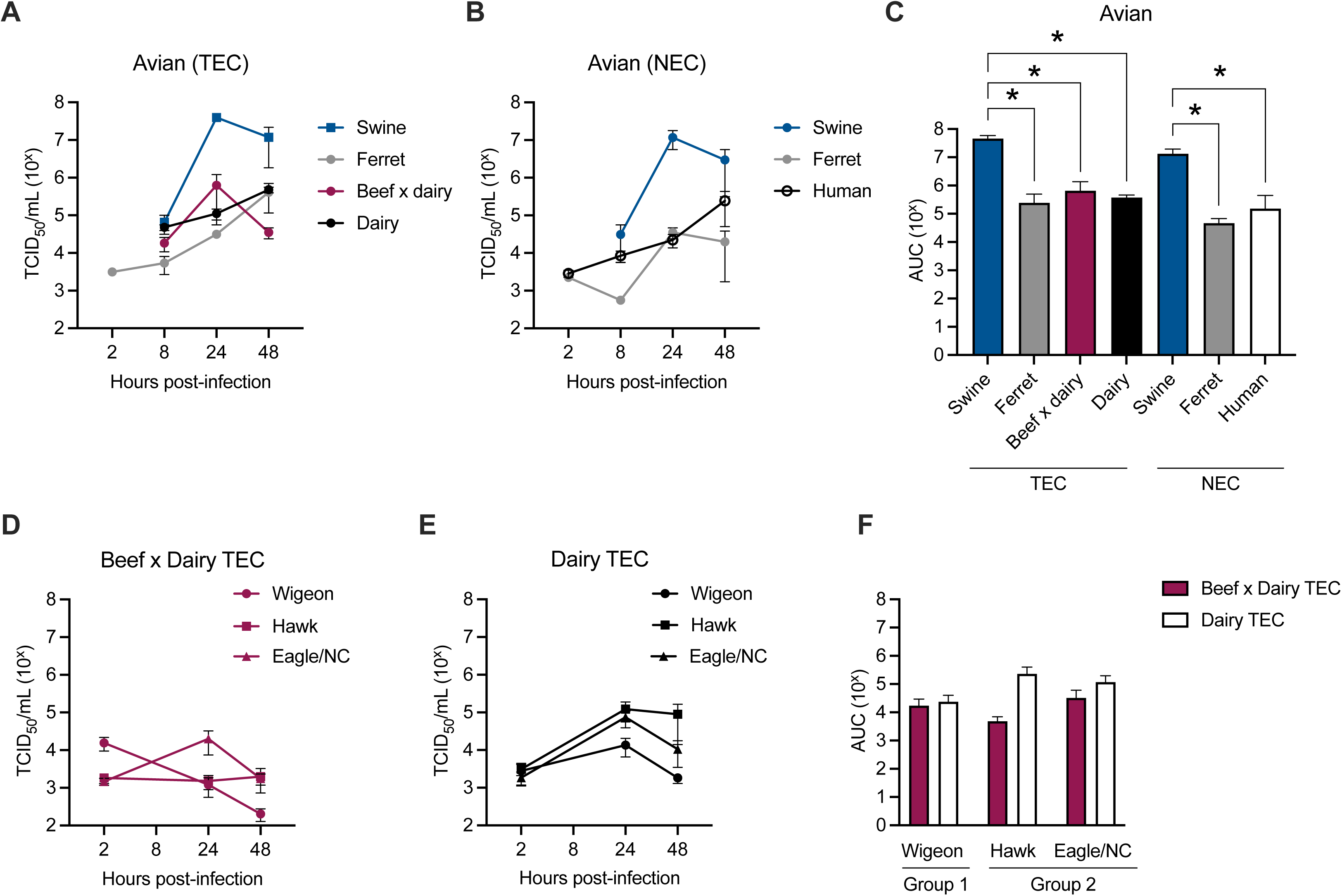
Avian derived clade 2.3.4.4b viruses do not replicate efficiently in respiratory epithelial cells. The indicated **(A)** TECs and **(B)** NECs were infected with A/bald eagle/Florida/W22-134-OP/2022 (H5N1) (eagle/FL) at an MOI of 0.5 and apical supernatants collected at the indicated times. Viral replication was measured by TCID_50_ assay. **(C)** Area under the curve (AUC) of (A-B). TEC: swine vs ferret *p*=0.0055; swine vs beef x dairy *p*=0.0058; swine vs dairy *p*=0.0056. NEC: swine vs ferret *p*=0.0053; swine vs human *p*=0.0029. Significance was determined by one-way ANOVA with multiple comparisons; \**p*<0.05, ^#^*p*<0.10. **(D)** Beef x dairy and **(E)** dairy TEC were infected with A/American wigeon/South Carolina/22-000345-001/2021 (H5N1) (wigeon); A/red-shouldered hawk/North Carolina/W22-121-2022 (H5N1) (hawk); or A/bald eagle/North Carolina/W22-140/2022 (H5N1) (eagle/NC) at a multiplicity of infection (MOI) of 0.5. Viral replication was measured as in (A-B). **(F)** Area under the curve of (D-E). Data represents 1 independent experiment, *n*=3-10 replicates per group. Error bars indicate standard error of the mean (SEM).

To ensure these findings were not unique to eagle/FL, we infected the bovine TECs with a panel of clade 2.3.4.4b avian derived viruses at an MOI of 0.5. This clade is subdivided into 2 groups based on genotype, and we chose strains representing both groups: group 1: A/American wigeon/South Carolina/22-000345-001/2021 (wigeon) and group 2: (A/red-shouldered hawk/North Carolina/W22-121/2022 (hawk) and A/bald eagle/North Carolina/W22-140/2022 (eagle/NC). Eagle/FL, used in Figures 4A-B, is part of group 1 (20). None of the avian viruses tested replicated in the bovine TEC, although titers of the group 2 viruses trended higher in the dairy TEC compared to beef x dairy (Figure 4D-F), This finding is supported by the underwhelming viral replication in the respiratory tract of cattle (5).

### Clade 2.3.4.4b viruses replicate in bovine mammary epithelial cells

A surprising feature of the current outbreak is the presence of high viral titers in milk, suggesting the mammary epithelium is infected (3, 6, 21). To test viral replication kinetics in this unique site, we isolated epithelial cells from the mammary gland tissues of the same cows that donated our TECs. Of note, the dairy cow was actively lactating at the time of cell isolation. Mammary epithelial cells (MEC) expressed pan-keratin confirming their identity as epithelial cells (9, 17), and could be differentiated in culture as evidenced by the expression of β-casein (14) (Figure 5). Like TECs, bovine and feline isolates could replicate to high titers in bovine MEC, although kinetics of the feline virus in beef x dairy MEC were slightly delayed (Figure 6A-B). Like the bovine TEC, the MEC were generally unable to support replication of A/Michigan/45/2015 (H1N1) or A/Switzerland/9715293/2013 (H3N2); however, the beef x dairy MEC were slightly permissive to H3N2 (Figure 6C). In contrast, bovine MEC supported the replication of the avian 2.3.4.4b viruses in almost all cases (Figure 6D-F) although we did note subtle differences between viral replication in the beef x dairy compared to the dairy cow, with replication trending higher in the dairy MEC (Figure 6F).

**Figure 5.**
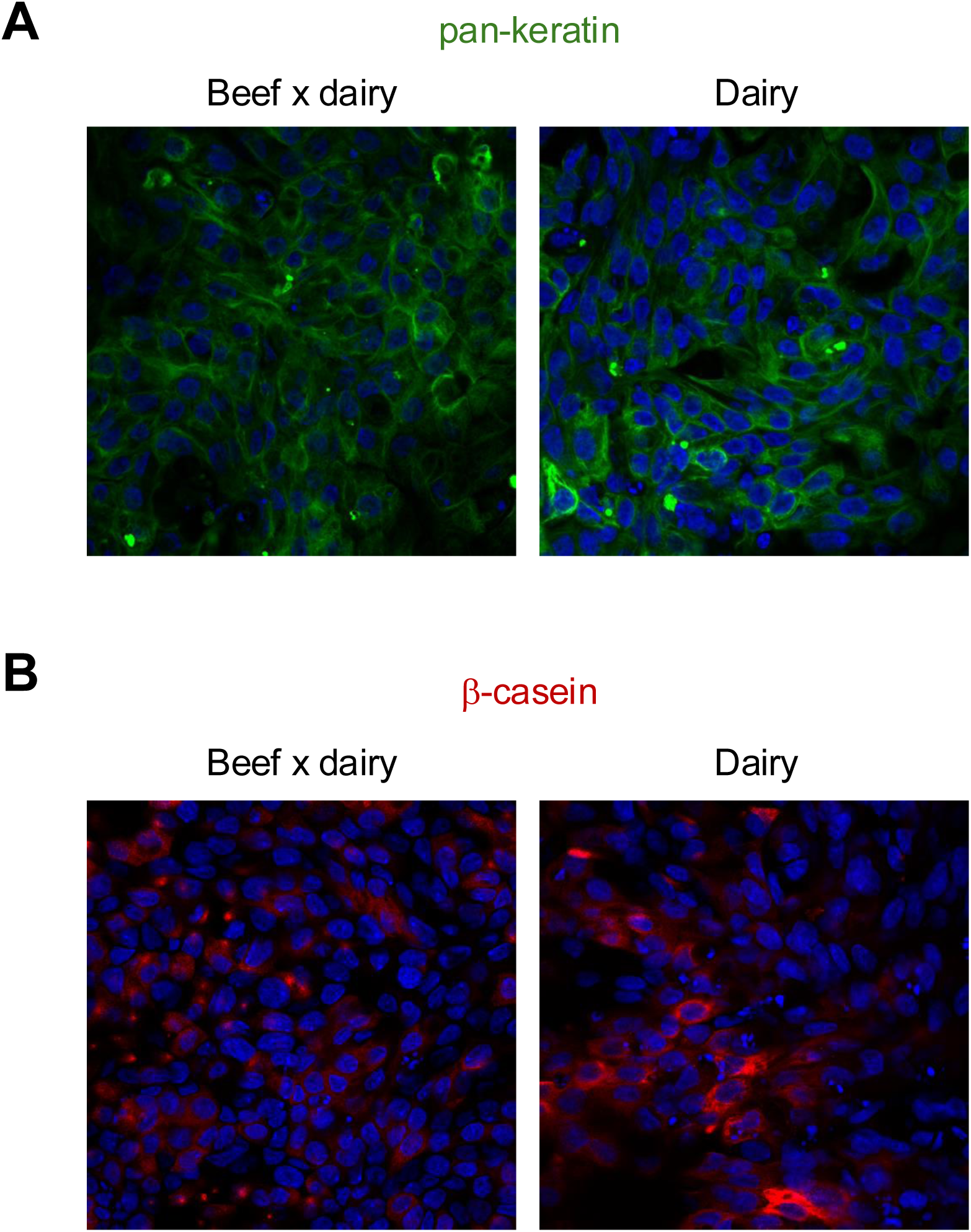
Differentiation of bovine mammary epithelial cells in culture. **(A)** Beef x dairy or **(B)** dairy mammary epithelial cells were fixed and immunostained for **(A)** pan-keratin (green), or **(B)** β-casein (red) and imaged by confocal microscopy. All samples were counterstained with DAPI (blue). Magnification is 40X. Data represents 2 independent experiments of n=2 per group.

**Figure 6.**
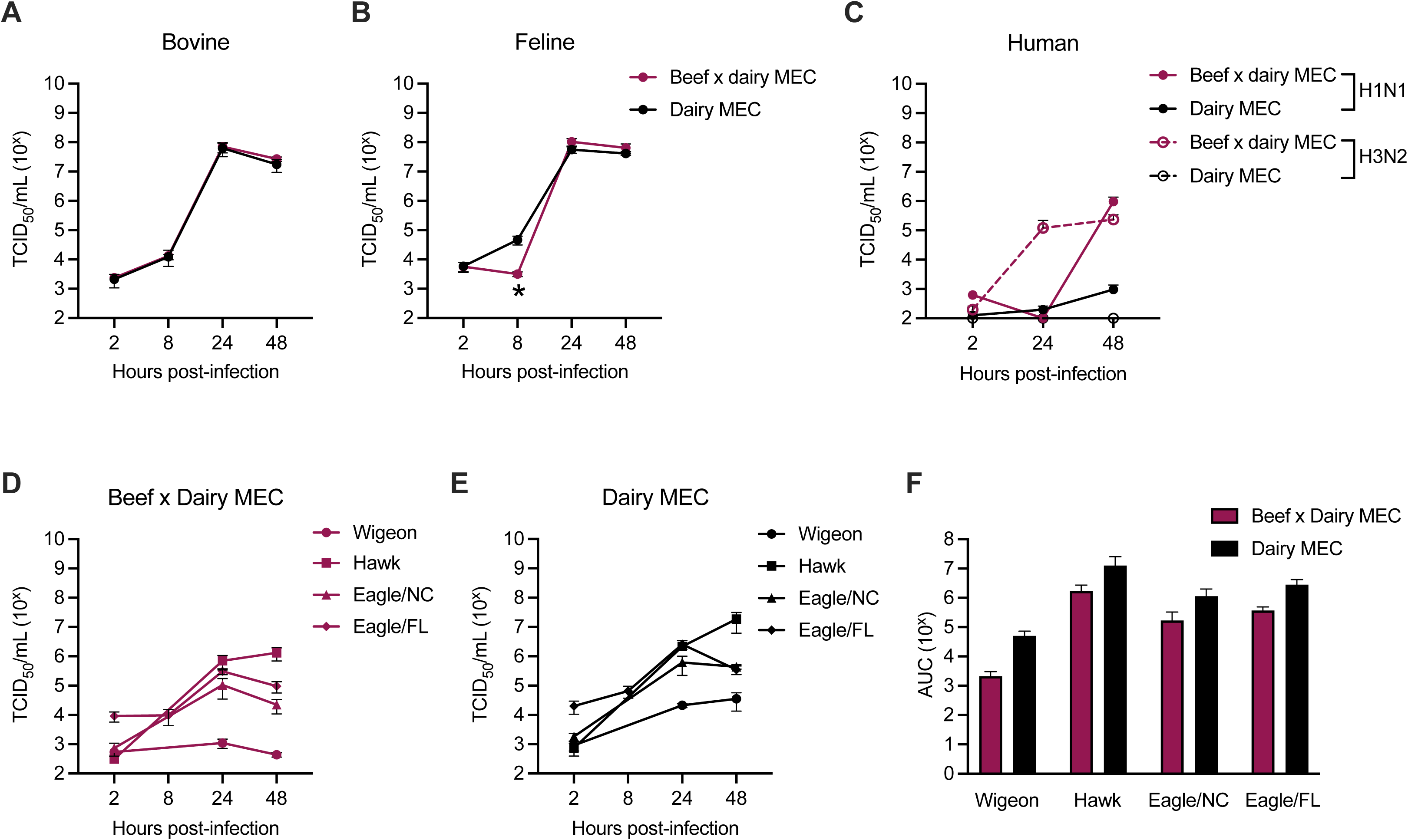
Replication of mammalian- and avian derived clade 2.3.4.4b viruses in bovine mammary epithelial cells. Mammary epithelial cells (MEC) from beef x dairy and dairy cows were infected with **(A)** A/bovine/Ohio/B24OSU-439/2024 (H5N1) (bovine) or **(B)** A/feline/New Mexico/2024 (H5N1) (feline) viruses at an MOI of 0.5. Apical supernatants were collected at the indicated time post-infection and viral replication was determined by TCID_50_ assay. Feline: *p*=0.0908 at 24 hours post-infection (hpi). **(C)** Cells were infected at an MOI of 0.5 with the seasonal human viruses A/Michigan/45/2015 (H1N1) or A/Switzerland/9715293/2013 (H3N2) as in (A-B). **(D)** Beef x dairy and **(E)** dairy MEC were infected with A/American wigeon/South Carolina/22-000345-001/2021 (H5N1), (wigeon); A/red-shouldered hawk/North Carolina/W22-121-2022 (H5N1), (hawk); A/bald eagle/North Carolina/W22-140/2022 (H5N1), (eagle/NC); or A/bald eagle/Florida/W22-134-OP/2022 (H5N1) (eagle/FL) at a multiplicity of infection (MOI) of 0.5. Viral titers were determined as in (A-C). **(F)** Area under the curve (AUC) of (D-E). Data represents 1 independent experiment, *n*=2-5 replicates per group. Error bars indicate standard error of the mean (SEM).

### Sialic acid receptor distribution is dependent on epithelial cell type and donor

Due to the differences in viral replication kinetics between beef x dairy and dairy epithelial cells, we asked whether sialic acid expression patterns differed between the two. We stained fully differentiated bovine TEC and MEC with the lectins MAA-II (α2,3) and SNA (α2,6) to assess sialic acid receptor expression. Interestingly, α2,3 expression was significantly increased in the dairy TEC compared to the beef x dairy TEC, suggesting avian viruses could be able to bind dairy TEC more efficiently (Figure 7A-C). Expression of α2,6 also trended higher in the dairy TEC, although the difference was not a striking (Figure 7B-C). Interestingly, expression of both α2,3 and α2,6 linkages was increased in the MEC compared to the TEC. However, MFI of both types of sialic acid receptors still trended towards higher expression in the dairy MEC than in the beef x dairy MEC (Figure 7D-F).

**Figure 7.**
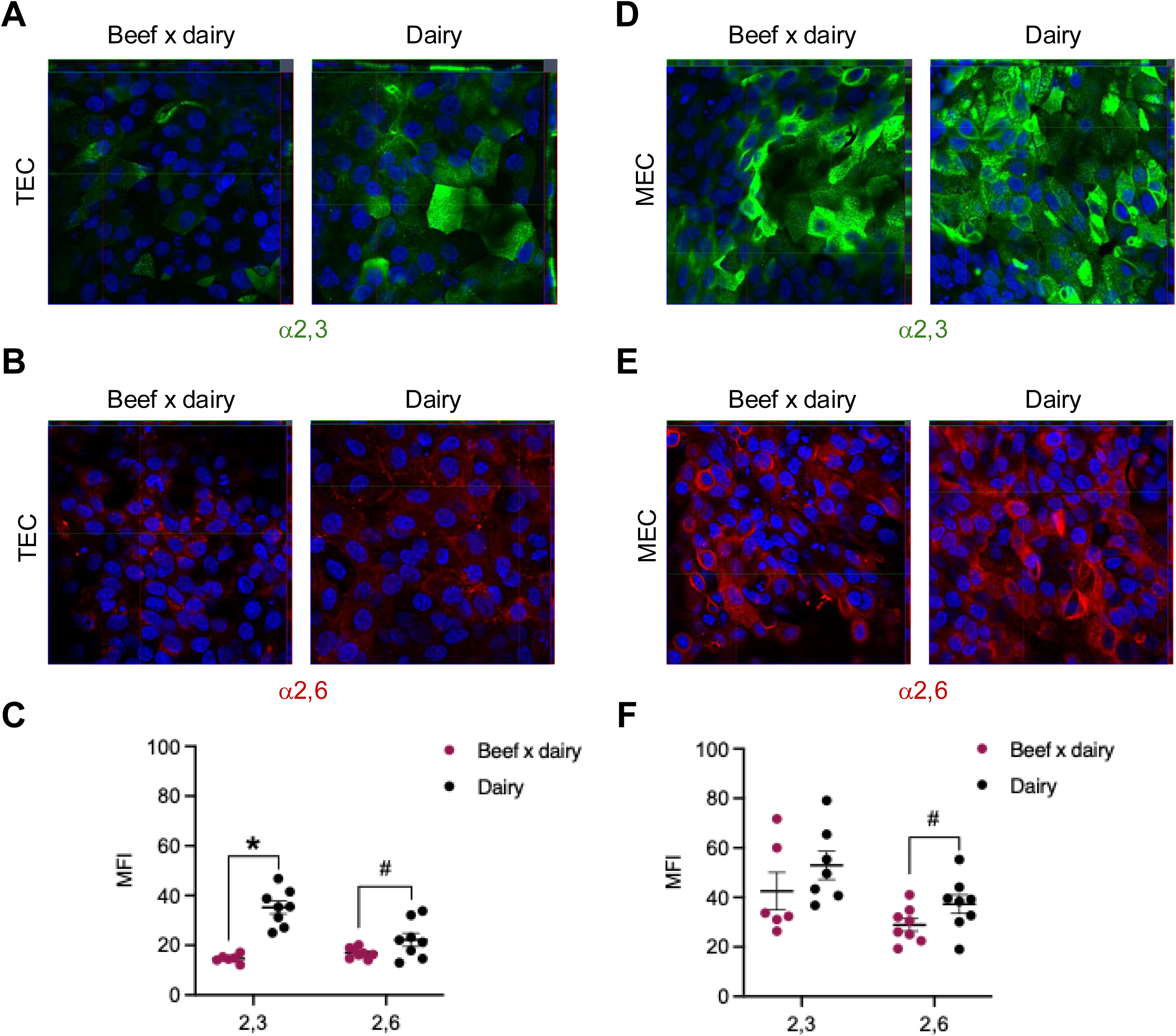
Sialic acid expression as determined by lectin staining in bovine tracheal and mammary epithelial cells. **(A)** Beef x dairy TEC and dairy TEC were cultured until fully differentiated. Cells were fixed and immunostained for α2,3 (MAA-II, green) and **(B)** α2,6 (SNA, red) sialic acid linkages and imaged by confocal microscopy. **(C)** Quantification of mean fluorescence intensity (MFI) shown in (A-B). *p*<0.0001 for α2,3 and *p*=0.0765 for α2,6. **(D-E)** Beef x dairy MEC and dairy MEC were stained as described in (A-B). **(F)** Quantification of mean fluorescence intensity (MFI) shown in (D-E). *p*=0.0831 for α2,6. Samples were counterstained with DAPI (blue). Magnification is 40X. Statistical significance was determined by unpaired *t* test; \**p*<0.05, ^#^*p*<0.10. Data represents 2 independent experiments of n=2 per group. Error bars represent standard error of the mean (SEM).

## Discussion

In this study, we developed a model of differentiated primary respiratory and mammary epithelial cells to better understand species and cellular tropism of the novel bovine H5N1 virus. We found that avian derived clade 2.3.4.4b viruses were unable to replicate well in respiratory epithelial cells of any type used in the study, except for swine. However, mammalian derived viruses could replicate to high titers in both TEC and NEC isolated from mammals. In contrast, bovine mammary epithelial cells could support replication of both avian and mammalian derived strains.

Avian influenza viruses are typically unable to infect mammals due to their preference for α2,3 sialic acid receptors, and mammalian respiratory epithelial cells primarily express α2,6 linkages in the upper respiratory tract (22–24). However, the mammalian derived bovine and feline viruses were able to replicate efficiently in TECs isolated from cow, swine, and ferret. There is data to support that some of the bovine strains can bind α2,6 linkages (2), but other studies have reported that the viruses do not (4, 25). This may be due to the usage of different strains of virus and different species from which the isolate was obtained among studies.

The nasal epithelium is traditionally the primary site of influenza infection in mammals. We therefore tested the bovine and feline isolates in NECs from swine, ferret, and human. Both isolates were able to replicate to high titers in all NECs tested. Overall, viral titers in swine NEC were significantly higher compared to ferret and human. This is troubling due to the propensity of influenza reassortment events in swine, and while there have previously been no reports of natural infection by clade 2.3.4.4b infections in swine (22), a strain was recently detected in swine from backyard farm in Oregon that had been in close contact with both poultry and humans (19).

We noted differences in viral kinetics between beef x dairy and dairy cow cells. For the bovine and the feline isolates, viral replication trended to be higher in the dairy TEC compared to beef x dairy TEC. Additionally, titers of the feline isolate were significantly higher in the dairy MEC at 24 hpi, suggesting replication may be delayed in beef x dairy MEC. While this could be due to donor-to-donor variation, one study showed increased binding of hemagglutinin (HA) protein in the mammary glands of lactating cattle, and little binding in a non-lactating cow (22). Another reported that there was not much variation in SNA lectin binding (targeting α2,6 receptors) in individual cows and cows of different breeds in the alveoli of the mammary gland, however there was wide variation in α2,6 expression in the ducts (16). Expression of both α2,3 and α2,6 receptors was increased on active alveoli (i.e. lactating) compared to inactive, leading the authors to hypothesize that lactating cattle may be more susceptible to infection (6, 16). This could also account for the high viral titers found in milk (3, 6). In accordance with these observations, our dairy cow was actively lactating at the time of euthanasia, while the beef x dairy cow was not. Inducing milk secretion in our MEC cultures may help shed light on this phenomenon in further studies.

While there have been no reports of human-to-human transmission thus far, differences in transmissibility of avian and mammalian derived clade 2.3.4.4b viruses imply the potential for adaptation. For example, a bovine strain isolated from a cow was unable to transmit via respiratory droplets in ferrets, while a bovine strain isolated from a human not only had the ability to transmit in ferrets, but could replicate *in vitro* at 33°C while the bovine strain could not (2, 26). There is some supporting evidence that the virus is adapting to transmit between mammals: cats on dairy farms that are fed raw milk have contracted the virus, but there have also been 2 cases reported where the infected cats had no contact with cattle, suggesting the virus may have transmitted by the cats eating wild birds (6, 21). Recently a wildlife sanctuary in Washington, U.S.A. lost 20 big cats (54% of their animals), including tigers, cougars, and servals, to the virus (27). Further, there have been recent reports of 2.3.4.4b viruses isolated from humans with severe disease that contain mutations associated with the ability to bind α2,6 (28–30), highlighting the need to understand viral tropism of mammalian derived viruses.

It is important to note that viral tropism is not completely dictated by sialic acid preferences. We detected α2,3 receptors on the cow TECs, particularly the dairy TEC, yet replication of avian derived 2.3.4.4b viruses was poor. In the case of cow MEC, α2,3 expression was high yet while avian 2.3.4.4b viruses could replicate in the MEC, the mammalian derived viruses were much more robust. One explanation for reduced replication of avian derived viruses could be lectin specificity. A study using recombinant HA protein to bind and identify tissue tropism of influenza virus revealed nonspecific binding of lectins to sialic acid receptors with similar linkages but are not receptors for influenza virus (22). Another explanation may be that we did whole organ isolation, and do not know if our MEC are alveolar epithelial cells, which express high levels of α2,3 and α2,6 which increase further during lactation, ductal epithelial cells that express lower levels (16), or a mix of both.

Understanding viral tropism is key to assess risk and evaluate pandemic potential of emerging influenza viruses. Here we show that nasal and tracheal epithelial cells from multiple species susceptible to influenza virus can support mammalian derived clade 2.3.4.4b H5N1 viruses, and bovine mammary cells are permissive to infection by both avian and mammalian viruses. These studies will provide information on factors that contribute to viral adaptation and transmission of avian viruses in the mammalian host.

## Methods

### Biosafety

All procedures involving infectious highly pathogenic avian influenza viruses were performed in the enhanced animal biosafety level 3 (ABSL3+) facility at St. Jude Children’s Research Hospital.

### Cells

Madin-Darby canine kidney (MDCK) cells (American Type Culture Collection, CCL-34) were cultured in MEM (Corning) containing GlutaMAX (2 mM; Gibco) and 10% fetal bovine serum (FBS) (HyClone). Human nasal epithelial cells (NEC) (abm) were cultured in Prigrow X.1 (abm). All animal cells were cultured in growth media [DMEM:F12 supplemented with penicillin (100 μg/mL), streptomycin (100 U/mL), GlutaMAX (2 mM), 5% FBS, bovine pituitary extract (30 μg/mL; Corning), cholera toxin (100 ng/mL; Sigma), insulin (10 μg/mL; Sigma), transferrin (5 μg/mL; Sigma), human epidermal growth factor (25 ng/mL; ThermoFisher), and retinoic acid (5 × 10^7^ M; Sigma)] as previously described (9). All cells were incubated at 37°C / 5% CO_2_.

### Isolation of bovine, ferret, and swine epithelial cells

Trachea and udder were obtained from 2 cows at the University of Wisconsin-Madison, courtesy of Dr. Sara Gragg. Swine trachea and nasal brushings were a gift of Dr. Henry Wan (Mississippi State University). Ferret tissues were harvested from male ferrets obtained from TripleF Farms (Elmyra, New York) under protocol 513 approved by the St. Jude Children’s Research Hospital Institutional Animal Care and Use Committee (IACUC).

*Tracheal epithelial cells*: excess connective tissue was removed and the trachea was cut into pieces (∼3 cm^2^) and placed into cold isolation media [DMEM:F12 (Corning) containing protease XIV (0.2%; Sigma), calcium acetate (16 μg/mL; Sigma), sodium acetate (16 μg/mL; Sigma), penicillin (100 μg/mL; Gibco), streptomycin (100 U/mL; Gibco), and amphotericin B (0.25 μg/mL; Gibco)] and incubated at 4°C overnight. The following morning, FBS was added to a final concentration of 10% to inactivate the protease. Epithelial cells were scraped from the inner surface of the trachea with a scalpel and collected by centrifugation at 200*g* for 5 minutes. The pellet was resuspended in isolation media containing 10% FBS and incubated in a 100 mm tissue culture treated dish at 37°C / 5% CO_2_ for 4 hours to allow any contaminating fibroblasts to adhere. The supernatant containing the epithelial cells was collected and centrifuged at 200*g* for 5 minutes.

#### Nasal epithelial cells

Nasal cell brushings from swine and ferret were collected with a FLOQSwab cytology brush (Copan Diagnostics). Swabs were washed with phosphate-buffered saline (PBS) to detach cells and centrifuged at 200*g* for 5 min. Both tracheal and nasal cells were resuspended in growth media and seeded onto tissue culture dishes pre-coated with rat tail collagen (0.05 μg/mL; Corning) diluted in 0.02 N glacial acetic acid. Bovine nasal brushings were unavailable.

#### Mammary epithelial cells

udders were sliced longitudinally to expose the mammary glands. Small (∼0.5 cm^3^) snips of tissue were collected and placed on 100 mm tissue culture treated dishes pre-coated with rat tail collagen. Growth media was added and the explants were incubated at 37°C / 5% CO_2_. Epithelial cells were harvested for further culture when they reached 80% confluence. For all epithelial cells, media was changed every 48 hours.

### Differentiation of primary epithelial cells

Tissue culture treated, 6.5 mm transwell inserts of 0.4 μm pore size, were coated with rat tail collagen (bovine, ferret, and swine cells) or extracellular matrix protein (abm) at manufacturer’s concentration (human nasal cells). Cells were seeded at a density of 3×10^4^/well. Respiratory cells were maintained in the corresponding growth media until the monolayer became 100% confluent at which point media was removed from the apical surface and the appropriate air-liquid interface (ALI) media was added to the basal compartment (Table 1). Mammary epithelial cell cultures remained submerged and were maintained in growth media throughout the differentiation process. Cells were considered fully differentiated once they developed cilia and secreted mucus (respiratory cells), or by the expression of β-casein (mammary cells).

**Table 1:**
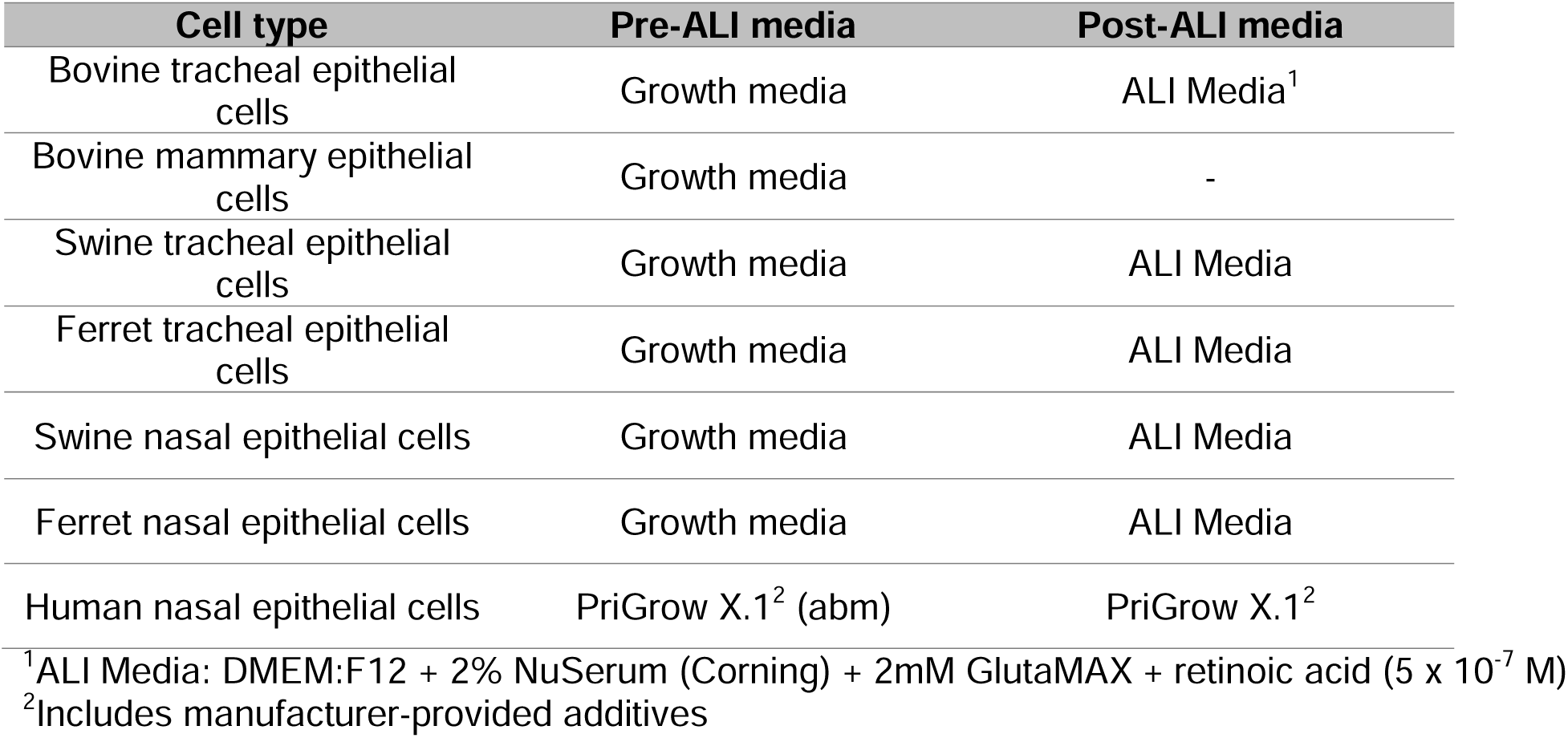
Appropriate growth and air-liquid interface (ALI) media by cell type.

### Viruses

H5N1 viruses were a kind gift from Dr. Richard Webby, St. Jude Children’s Research Hospital (A/red-shouldered hawk/North Carolina/W22-121/2022, A/American wigeon/South Carolina/22-000345-001/2021, A/bald eagle/North Carolina/W22-140/2022, A/bald eagle/Florida/W22-134-OP/2022, A/bovine/Ohio/B24OSU-439/2024) and Dr. Kiril Dimitrov, Texas A&M (A/feline/New Mexico/2024). Viruses were propagated through 9-day old embryonated chicken eggs as previously described (31). A/Michigan/45/2015 (H1N1) and A/Switzerland/9715293/2013 (H3N2) were propagated in MDCK as previously described (32). Viral titers were measured by tissue culture infectious dose 50 (TCID_50_) assay.

### Influenza infections

To infect MDCK, cells were seeded into a 24-well tissue culture plate at a density of 5 × 10^5^ per well. The monolayer was washed twice with PBS and viral inoculum diluted in infection media (MEM supplemented with 0.075% bovine serum albumin and 2 mM GlutaMAX) at a multiplicity of infection (MOI) of 0.01 or 0.001. Plates were incubated at 37°C / 5% CO_2_ for 1 hour before removing viral inoculum. The surface of the monolayer was washed twice with PBS to remove any unbound virus and 1 mL of infection media added. Supernatants were collected at the indicated times and infectious virus was measured by TCID_50_ assay.

For infections of differentiated primary cells, the apical surface was washed twice with PBS. Viral inoculum diluted in infection media was added to the apical surface and plates were incubated for 1 hour. Viral inoculum was removed and the apical surface washed twice with PBS. To collect supernatants, infection media was added to the apical compartment and plates were incubated for 30 min at 37°C / 5% CO_2_. Supernatants were collected and infectious virus was measured by TCID_50_ assay.

### Tissue culture infectious dose 50 (TCID_50_) assay

MDCK were seeded into 96-well cell culture plates at a density of 2.5 × 10^4^ per well and incubated overnight at 37°C / 5% CO_2_. The next day, cells were washed twice with PBS and inoculated with ten-fold serial dilutions of sample diluted in infection media. Plates were returned to the incubator for 72 hours. Following, 50 μL of cell supernatant were transferred to a V-bottom plate and 0.5% packed turkey red blood cells diluted in PBS were added (50 μL). Plates were incubated for 30 minutes at room temperature and viral titers determined via the presence or absence of hemagglutination. Titers were calculated according to the method of Reed and Muench (33).

### Immunofluorescent staining

Differentiated cells were fixed in 4% paraformaldehyde (Fisher Scientific) diluted in PBS for 20 min. Cells were washed 3X with PBS and permeabilized with 0.1% TritonX-100 (Sigma) in PBS for 15 min. Cells were washed again and blocked in 5% bovine serum albumin fraction V (HyClone) diluted in PBS+0.05% Tween-20 (Fisher Scientific) (PBST) at room temperature for 1 h. Block was removed and replaced with primary antibody [Sambucus nigra (SNA)-FITC (Vector Laboratories; 15 μg/mL), biotinylated Maackia amurensis (MAA)-II (Vector Laboratories; 15 μg/mL), anti-β-casein (ThermoFisher; 10 μg/mL), anti-MUC1 (Abcam clone EP1024Y; 0.25 μg/mL), anti-β-tubulin (Biolegend clone TU27; 2 μg/mL), or anti-pan-keratin conjugated to AlexaFluor 488 (Biolegend clone C-11; 5 μg/mL)] diluted in 1% BSA in PBST and incubated overnight at 4°C. After washing 3X with PBST, cells were incubated with the appropriate secondary antibody (streptavidin-TexasRed (Vector Laboratories; 15 μg/mL), anti-rabbit AlexaFluor 555 (Invitrogen, 10 μg/mL), or anti-rabbit AlexaFluor 488 (Invitrogen, 10 μg/mL)) and counterstained with Hoescht dye (1 μg/mL) diluted in 1% BSA PBST and incubated for 1 h at room temperature. The transwell membrane was excised with a scalpel and mounted onto a glass slide with ProLong Gold AntiFade (Invitrogen) and cured for 24h before imaging. Images were captured with a Zeiss LSM 780 Observer.Z1. Data was analyzed with Zen Black 2012 SP 5 (14.0.28.201).

### Statistical analysis

Statistical analysis was performed as described in figure legends by GraphPad Prism version 10.3.1. Significance was defined as *p*<0.05 (denoted by an asterisk), and values of 0.05<*p*<0.1 were considered trending (denoted by # symbol).

## Acknowledgements

The authors would like to thank David Carey, Heather Weinberg, and the ABSL3+ staff at the Animal Resource Center at St. Jude Children’s Research Hospital. We thank the St. Jude Children’s Research Hospital Cell and Tissue Imaging Core facility. We thank Dr. Richard Webby for the bovine H5N1 influenza strain and Dr. Henry Wan at Mississippi State University for providing swine tissues. Finally, we extend our thanks for all the cows, ferrets, swine, and anonymous human donors for their contribution to science. Funding was provided by the National Institutes of Health Centers for Excellence in Influenza Research and Surveillance (CEIRS) contact no. HHSN27220140006C (SSC), National Institute of Health, Institutional Postdoctoral Training Grants (T32) Infectious Disease Therapeutics T32AI106700-8 (PHB and TR), the National Institute of Health Ruth L. Kirschstein Postdoctoral Individual National Research Service Award F32AI183804 (PHB). and the American Lebanese Syrian Associated Charities (ALSAC) (SSC).

## Author contributions

Conceptualization: VM, SL, SSC; Data curation: VM; Formal analysis: VM, MS; Funding acquisition: SSC; Investigation: VM, SC, MS, BS, PHB, ER, BL, DM, TR, PB; Methodology: VM, SC, SSC; Project administration: VM, SSC; Resources: SG, KD, SL, SSC; Software: N/A; Supervision: SL, SSC; Validation: VM, SC, MS, PB; Visualization: VM, MS, SSC; Writing – original draft: VM, SSC; Writing – review and editing: SSC, VM, DRM, SL, PHB.

## Supplemental material

**Supplemental Figure 1.**
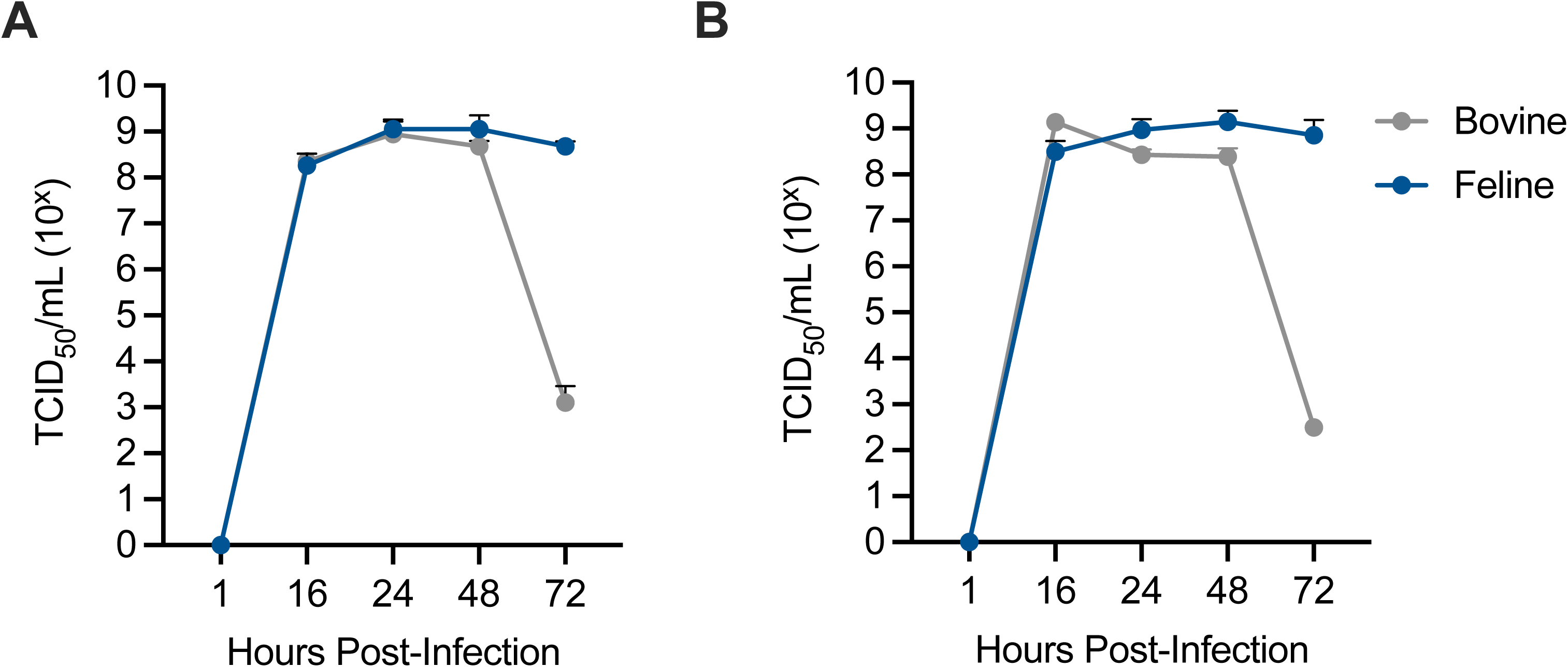
Viral growth kinetics of bovine and feline H5N1 *in vitro*. MDCK cells were infected with A/bovine/Ohio/B24OSU-439/2024 (H5N1) or A/feline/New Mexico/2024 (H5N1) at a multiplicity of infection (MOI) of **(A)** 0.001 or **(B)** 0.01. Supernatants were collected at the indicated time. Data represents 1 independent experiment of *n*=3-6 per group. Error bars represent standard deviation (SD).

## Notes

### Competing Interest Statement

The authors have declared no competing interest.

